# Fast and robust detection of colistin resistance in *Escherichia coli* using the MALDI Biotyper Sirius mass spectrometry system

**DOI:** 10.1101/752600

**Authors:** R. Christopher D. Furniss, Laurent Dortet, William Bolland, Oliver Drews, Katrin Sparbier, Rémy A. Bonnin, Alain Filloux, Markus Kostrzewa, Despoina A.I. Mavridou, Gerald Larrouy-Maumus

**Affiliations:** MRC Centre for Molecular Bacteriology and Infection, Department of Life Sciences, Faculty of Natural Sciences, Imperial College London, London, SW7 2AZ, UK; Department of Bacteriology-Hygiene, Bicêtre Hospital, Assistance Publique - Hôpitaux de Paris, Le Kremlin-Bicêtre, France; EA7361 “Structure, dynamic, function and expression of broad spectrum β-lactamases”, Paris-Sud University, LabEx Lermit, Faculty of Medecine, Le Kremlin-Bicêtre, France; French National Reference Centre for Antibiotic Resistance, France; Bruker Daltonik GmbH, Bremen, Germany

## Abstract

Polymyxin antibiotics are a last-line treatment for multidrug-resistant Gram-negative bacteria. However, the emergence of colistin resistance, including the spread of mobile *mcr* genes, necessitates the development of improved diagnostics for the detection of colistin-resistant organisms in hospital settings. The recently developed MALDIxin test enables detection of colistin resistance by MALDI-TOF mass spectrometry in less than 15 minutes but is not optimized for the mass spectrometers commonly found in clinical microbiology laboratories. In this study, we adapted the MALDIxin test for the MALDI Biotyper Sirius MALDI-TOF mass spectrometry system (Bruker Daltonics). We optimized the sample preparation protocol using a set of 6 MCR-expressing *Escherichia coli* clones and validated the assay with a collection of 40 *E. coli* clinical isolates, including 19 MCR producers, 12 chromosomally-resistant isolates and 9 polymyxin-susceptible isolates. We calculated Polymyxin resistance ratio (PRR) values from the acquired spectra; a PRR value of zero, indicating polymyxin susceptibility, was obtained for all colistin-susceptible *E. coli* isolates, whereas positive PRR values, indicating resistance to polymyxins, were obtained for all resistant strains independent of the genetic basis of resistance. Thus, we report a preliminary feasibility study showing that an optimized version of the MALDIxin test, adapted for the routine MALDI Biotyper Sirius, provides an unbiased, fast, reliable, cost-effective and high-throughput way of detecting colistin resistance in clinical *E. coli* isolates.

## INTRODUCTION

Antibiotic resistance is an issue of global importance and one of the defining public health concerns of our time (1). The limited pipeline of novel antimicrobials and the spread of multidrug-resistant (MDR) organisms have increased our reliance on a few last-line antibiotics for the treatment of MDR Gram-negative bacteria. Chief amongst these last-resort agents are the polymyxin antibiotics, polymyxin B and colistin (2, 3).

In Gram-negative bacteria like *Escherichia coli*, polymyxin resistance mostly occurs as a consequence of lipopolysaccharide (LPS) modifications, in the form of addition of the cationic groups phosphoethanolamine (pETN) and/or 4-amino-L-arabinose (L-Ara4N) to the Lipid A portion of LPS (4, 5). These Lipid A modifications often arise due to alterations to the PmrAB and PhoPQ two-component systems, mutations to the negative regulator of PhoPQ, MgrB, or because of the activity of plasmid-borne pETN transferases called mobile colistin resistance (MCR) enzymes (6). The first MCR enzyme, MCR-1, was reported in 2016 (7) and this discovery was followed by the rapid identification of other mobile polymyxin resistance genes. To date a further eight MCR proteins have been described. These enzymes cluster into four main groups: MCR-1-like (MCR-1, −2, −6), MCR-3-like (MCR-3, −7, −8, −9), MCR-4-like (MCR-4) and MCR-5-like (MCR-5) (8–11).

Detection of colistin resistance currently relies on minimum inhibitory concentration (MIC) determination using broth microdilution (BMD), a slow process which, despite being the gold standard for polymyxin susceptibility testing, has been subject to reliability and standardization problems (6, 12). Additionally, routine detection of colistin resistance by conventional methods such as polymerase-chain-reaction (PCR)-based testing is challenging due to the wide range of chromosomal mutations which can give rise to colistin resistance (6) and the low sequence identity of the *mcr* genes (using *mcr-1* as a reference: *mcr-2,* 77.6%; *mcr-3,* 49.2%; *mcr-4,* 46.8%; *mcr-5,* 50. 5%; *mcr-6,* 78.3%; *mcr-7,* 49.9%; *mcr-8,* 47.8%; *mcr-9,* 57.69%). This means that PCR-based detection methods are insensitive to all but the best-characterized chromosomal mutations and to the emergence of new *mcr* genes. Therefore, there is an urgent need to develop a fast, robust and high-throughput assay, accessible to all diagnostic microbiology laboratories, that uses an unbiased approach to detect colistin resistance arising from both known and novel chromosomal mutations or MCR proteins.

Recently we developed the MALDIxin test, a diagnostic tool based on Matrix-assisted laser desorption/ionization-time of flight (MALDI-TOF) mass spectrometry that can be used to detect colistin resistance using intact bacteria in less than 15 minutes (13, 14). Although fast and effective, this test was not optimized for routine use in diagnostic microbiology laboratories, the main limitation being that it was not developed for the MALDI-TOF mass spectrometers widely used for bacterial identification in these settings. More specifically, our previous studies were performed on a research instrument operating in the high-resolution reflector mode, whilst MALDI-TOF systems in clinical microbiology laboratories employ lower resolution linear mode measurements. Here, we report a preliminary feasibility study showing that an optimized version of the MALDIxin test, designed for the low-resolution linear mode employed by the MALDI Biotyper Sirius system (Bruker Daltonics), accurately identifies colistin resistance in clinical *E. coli* isolates irrespective of its genetic basis by detecting addition of both pETN and L-Ara4N moieties to Lipid A.

## MATERIALS AND METHODS

### Bacterial strains

For the construction of MCR-producing *E. coli* clones (Table 1), *mcr* variants were cloned into pDM1 (GenBank MN128719), an isopropyl β-D-1-thiogalactopyranoside (IPTG)-inducible derivative of pACYC184; protein expression from this vector is only induced after addition of IPTG to the culture media. For *mcr-1, mcr-2, mcr-4, mcr-5* and *mcr-8* the SacI/XmaI sites of the vector were used, whilst for *mcr-3,* the NdeI/XmaI sites were used. A collection of 40 *E. coli* clinical isolates (Table 1), including 19 MCR producers, 12 chromosomally-resistant isolates and 9 colistin susceptible isolates, was used for validation of the MALDIxin test.

**Table 1.**
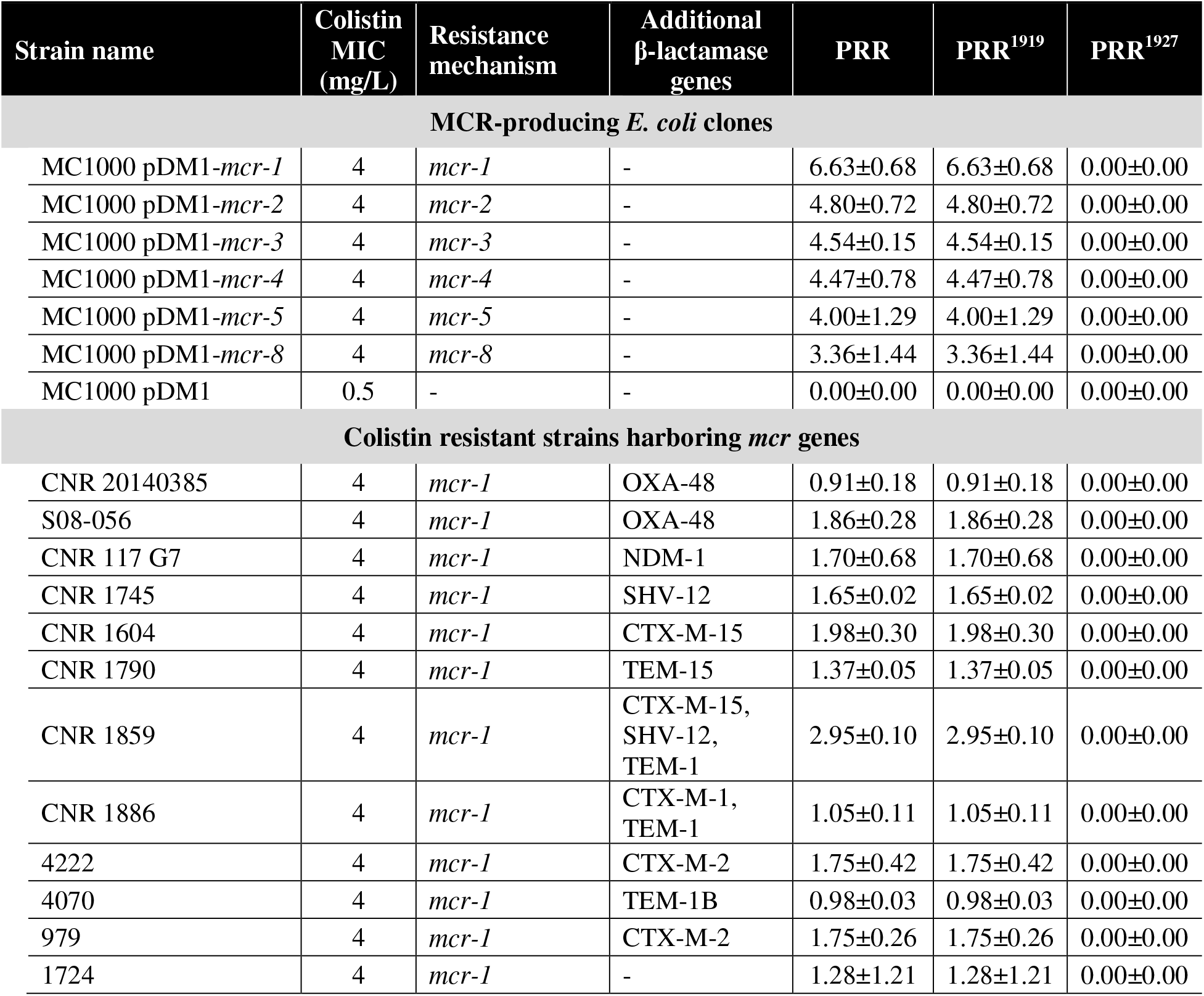
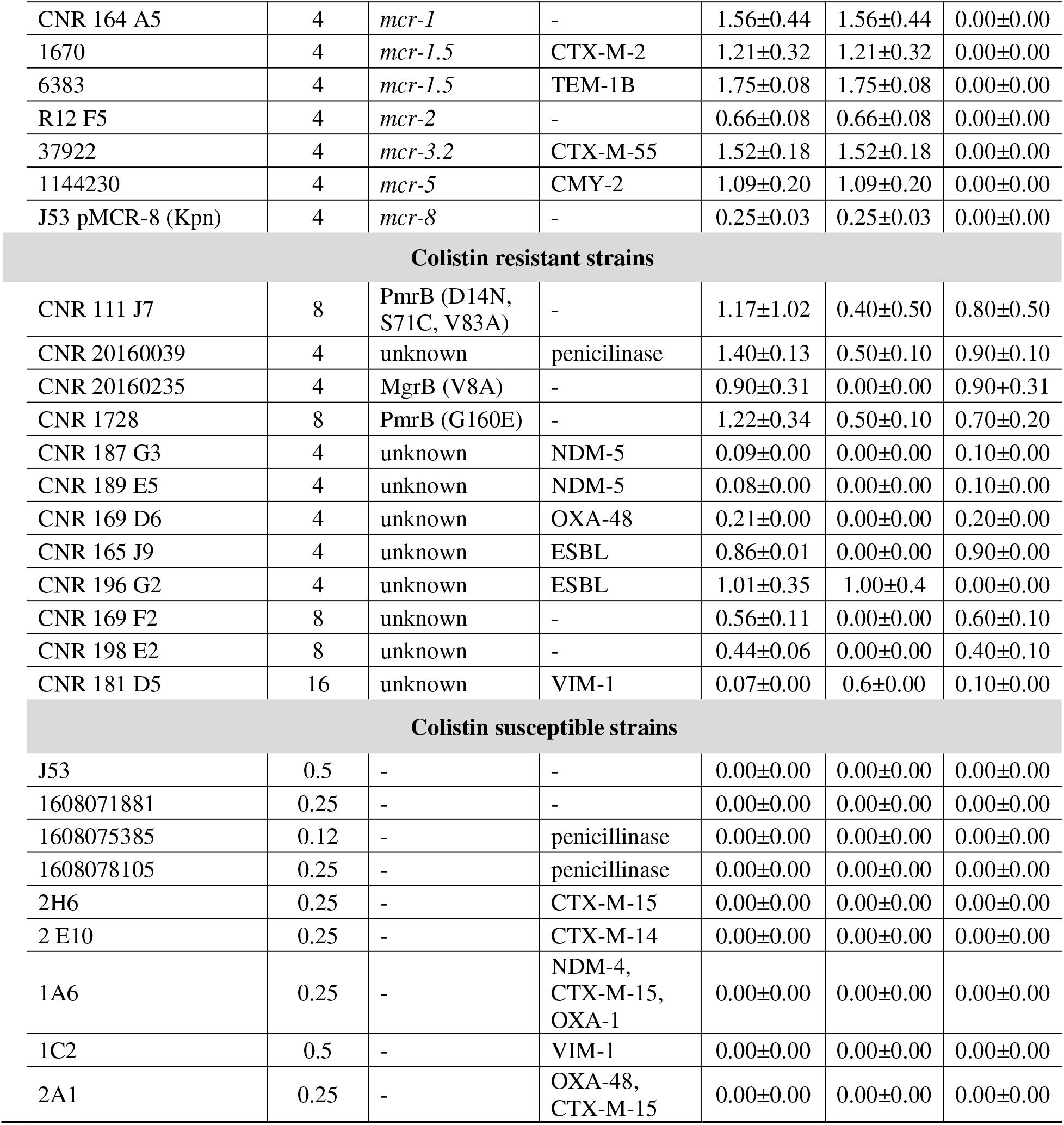
PRR values for the MCR-producing *E. coli* clones, colistin-resistant clinical *E. coli* strains and susceptible *E. coli* strains used in this study. PRR is calculated by summing the intensities of the Lipid A peaks attributed to the addition of pETN (*m/z* 1919.2) and L-Ara4N (*m/z* 1927.2) and dividing this number by the intensity of the peak corresponding to native Lipid A (*m/z* 1796.2); PRR = (1919.2 intensity+1927.2 intensity) / 1796.2 intensity. PRR^1919^ and PRR^1927^ indicate the contribution of specific Lipid A modification(s) (pETN and/or L-Ara4N) to the overall PRR value. PRR^1919^ and PRR^1927^ are calculated by dividing the intensity of the peak at the appropriate *m/z* (*m/z* 1919.2 and *m/z* 1927.2 for pETN and L-Ara4N addition, respectively) by the intensity of the peak corresponding to native Lipid A (*m/z* 1796.2).

| Strain name | Colistin MIC (mg/L) | Resistance mechanism | Additional β-lactamase genes | PRR | $PRR^{1919}$ | $PRR^{1927}$ |
| --- | --- | --- | --- | --- | --- | --- |
| **MCR-producing *E. coli* clones** |  |  |  |  |  |  |
| MC1000 pDM1-*mcr-1* | 4 | *mcr-1* | - | 6.63±0.68 | 6.63±0.68 | 0.00±0.00 |
| MC1000 pDM1-*mcr-2* | 4 | *mcr-2* | - | 4.80±0.72 | 4.80±0.72 | 0.00±0.00 |
| MC1000 pDM1-*mcr-3* | 4 | *mcr-3* | - | 4.54±0.15 | 4.54±0.15 | 0.00±0.00 |
| MC1000 pDM1-*mcr-4* | 4 | *mcr-4* | - | 4.47±0.78 | 4.47±0.78 | 0.00±0.00 |
| MC1000 pDM1-*mcr-5* | 4 | *mcr-5* | - | 4.00±1.29 | 4.00±1.29 | 0.00±0.00 |
| MC1000 pDM1-*mcr-8* | 4 | *mcr-8* | - | 3.36±1.44 | 3.36±1.44 | 0.00±0.00 |
| MC1000 pDM1 | 0.5 | - | - | 0.00±0.00 | 0.00±0.00 | 0.00±0.00 |
| **Colistin resistant strains harboring *mcr* genes** |  |  |  |  |  |  |
| CNR 20140385 | 4 | *mcr-1* | OXA-48 | 0.91±0.18 | 0.91±0.18 | 0.00±0.00 |
| S08-056 | 4 | *mcr-1* | OXA-48 | 1.86±0.28 | 1.86±0.28 | 0.00±0.00 |
| CNR 117 G7 | 4 | *mcr-1* | NDM-1 | 1.70±0.68 | 1.70±0.68 | 0.00±0.00 |
| CNR 1745 | 4 | *mcr-1* | SHV-12 | 1.65±0.02 | 1.65±0.02 | 0.00±0.00 |
| CNR 1604 | 4 | *mcr-1* | CTX-M-15 | 1.98±0.30 | 1.98±0.30 | 0.00±0.00 |
| CNR 1790 | 4 | *mcr-1* | TEM-15 | 1.37±0.05 | 1.37±0.05 | 0.00±0.00 |
| CNR 1859 | 4 | *mcr-1* | CTX-M-15, SHV-12, TEM-1 | 2.95±0.10 | 2.95±0.10 | 0.00±0.00 |
| CNR 1886 | 4 | *mcr-1* | CTX-M-1, TEM-1 | 1.05±0.11 | 1.05±0.11 | 0.00±0.00 |
| 4222 | 4 | *mcr-1* | CTX-M-2 | 1.75±0.42 | 1.75±0.42 | 0.00±0.00 |
| 4070 | 4 | *mcr-1* | TEM-1B | 0.98±0.03 | 0.98±0.03 | 0.00±0.00 |
| 979 | 4 | *mcr-1* | CTX-M-2 | 1.75±0.26 | 1.75±0.26 | 0.00±0.00 |
| 1724 | 4 | *mcr-1* | - | 1.28±1.21 | 1.28±1.21 | 0.00±0.00 |
| CNR 164 A5 | 4 | *mcr-1* | - | 1.56±0.44 | 1.56±0.44 | 0.00±0.00 |
| 1670 | 4 | *mcr-1.5* | CTX-M-2 | 1.21±0.32 | 1.21±0.32 | 0.00±0.00 |
| 6383 | 4 | *mcr-1.5* | TEM-1B | 1.75±0.08 | 1.75±0.08 | 0.00±0.00 |
| R12 F5 | 4 | *mcr-2* | - | 0.66±0.08 | 0.66±0.08 | 0.00±0.00 |
| 37922 | 4 | *mcr-3.2* | CTX-M-55 | 1.52±0.18 | 1.52±0.18 | 0.00±0.00 |
| 1144230 | 4 | *mcr-5* | CMY-2 | 1.09±0.20 | 1.09±0.20 | 0.00±0.00 |
| J53 pMCR-8 (Kpn) | 4 | *mcr-8* | - | 0.25±0.03 | 0.25±0.03 | 0.00±0.00 |
| Colistin resistant strains |  |  |  |  |  |  |
| CNR 111 J7 | 8 | PmrB (D14N, S71C, V83A) | - | 1.17±1.02 | 0.40±0.50 | 0.80±0.50 |
| CNR 20160039 | 4 | unknown | penicillinase | 1.40±0.13 | 0.50±0.10 | 0.90±0.10 |
| CNR 20160235 | 4 | MgrB (V8A) | - | 0.90±0.31 | 0.00±0.00 | 0.90±0.31 |
| CNR 1728 | 8 | PmrB (G160E) | - | 1.22±0.34 | 0.50±0.10 | 0.70±0.20 |
| CNR 187 G3 | 4 | unknown | NDM-5 | 0.09±0.00 | 0.00±0.00 | 0.10±0.00 |
| CNR 189 E5 | 4 | unknown | NDM-5 | 0.08±0.00 | 0.00±0.00 | 0.10±0.00 |
| CNR 169 D6 | 4 | unknown | OXA-48 | 0.21±0.00 | 0.00±0.00 | 0.20±0.00 |
| CNR 165 J9 | 4 | unknown | ESBL | 0.86±0.01 | 0.00±0.00 | 0.90±0.00 |
| CNR 196 G2 | 4 | unknown | ESBL | 1.01±0.35 | 1.00±0.4 | 0.00±0.00 |
| CNR 169 F2 | 8 | unknown | - | 0.56±0.11 | 0.00±0.00 | 0.60±0.10 |
| CNR 198 E2 | 8 | unknown | - | 0.44±0.06 | 0.00±0.00 | 0.40±0.10 |
| CNR 181 D5 | 16 | unknown | VIM-1 | 0.07±0.00 | 0.6±0.00 | 0.10±0.00 |
| Colistin susceptible strains |  |  |  |  |  |  |
| J53 | 0.5 | - | - | 0.00±0.00 | 0.00±0.00 | 0.00±0.00 |
| 1608071881 | 0.25 | - | - | 0.00±0.00 | 0.00±0.00 | 0.00±0.00 |
| 1608075385 | 0.12 | - | penicillinase | 0.00±0.00 | 0.00±0.00 | 0.00±0.00 |
| 1608078105 | 0.25 | - | penicillinase | 0.00±0.00 | 0.00±0.00 | 0.00±0.00 |
| 2H6 | 0.25 | - | CTX-M-15 | 0.00±0.00 | 0.00±0.00 | 0.00±0.00 |
| 2 E10 | 0.25 | - | CTX-M-14 | 0.00±0.00 | 0.00±0.00 | 0.00±0.00 |
| 1A6 | 0.25 | - | NDM-4,<br>CTX-M-15,<br>OXA-1 | 0.00±0.00 | 0.00±0.00 | 0.00±0.00 |
| 1C2 | 0.5 | - | VIM-1 | 0.00±0.00 | 0.00±0.00 | 0.00±0.00 |
| 2A1 | 0.25 | - | OXA-48,<br>CTX-M-15 | 0.00±0.00 | 0.00±0.00 | 0.00±0.00 |

### Genotype determination

PCR-based amplification and DNA sequencing was used to determine the genotypes of clinical isolates (Table 1), as necessary. Identification of *mcr* genes was performed by multiplex PCR as previously described (15) and β-lactamases genes were identified using in-house multiplex PCR protocols.

### Susceptibility testing

Colistin MICs for clinical isolates were manually determined using BMD, according to the Clinical and Laboratory Standards Institute (CLSI) and the European Committee on Antimicrobial Susceptibility Testing (EUCAST) guidelines. As such, cation-adjusted Mueller-Hinton broth was used in conjunction with plain polystyrene laboratory consumables and the sulfate salt of colistin. No additives were used at any stage of the testing process. For the laboratory *E. coli* clones, which were only used for protocol optimization, 0.5 mM IPTG was added to the BMD growth media to induce expression of the MCR enzymes. The colistin MIC of all tested strains was determined three times and was found to be identical at each repeat. Results were interpreted using EUCAST breakpoints as updated in 2018 (http://www.eucast.org/clinical_breakpoints/).

### Optimized MALDIxin test for the MALDI Bioptyper Sirius

A 10 μL inoculation loop of bacteria, grown on Mueller-Hinton agar for 18-24 hours, was resuspended in 200 μL of water. Mild-acid hydrolysis was performed on 100 μL of this suspension, by adding 100 μl of 2 v/v % acetic acid and incubating the mixture at 98°C for 10 min. Hydrolyzed cells were centrifuged at 17,000 *x g* for 2 min, the supernatant was discarded and the pellet was resuspended in ultrapure water to a density of McFarland 10. 0.4 μL of this suspension was loaded onto the target and immediately overlaid with 1.2 μL of a matrix consisting of a 9:1 mixture of 2,5-dihydroxybenzoic acid and 2-hydroxy-5-methoxybenzoic acid (super-DHB) (Sigma Aldrich) dissolved in 90/10 v/v chloroform/methanol to a final concentration of 10 mg/mL. The bacterial suspension and matrix were mixed directly on the target by pipetting and the mix dried gently under a stream of air for less than one minute. MALDI-TOF mass spectrometry analysis was performed with a MALDI Biotyper Sirius, (Bruker Daltonics) using the newly introduced linear negative-ion mode.

### Data analysis

Manual peak picking at masses relevant to colistin resistance was performed on the obtained mass spectra and the corresponding signal intensities at these defined masses was determined. The sum of the intensities of the Lipid A peaks attributed to addition of pETN (*m/z* 1919.2) and L-Ara4N (*m/z* 1927.2) was divided by the intensity of the peak corresponding to native Lipid A (*m/z* 1796.2). The resulting value is termed the polymyxin resistance ratio (PRR). A PRR of zero indicates colistin susceptibility, whilst a positive value indicates colistin resistance. PRR^1919^ values were determined by dividing the intensity of the peak at *m/z* 1919 alone by the native Lipid A peak and PRR^1927^ values were determined by dividing the intensity of the peak at *m/z* 1927 alone by the native Lipid A peak. All mass spectra were generated and analyzed in technical triplicate (i.e. measurements of each sample were repeated three times) and biological triplicate (i.e. the entire experiment was repeated on three separate days using separately grown bacteria and separate materials).

## RESULTS

To allow the use of the MALDIxin test on the MALDI Biotyper Sirius system it was necessary to optimize the sample preparation protocol. This optimization was carried out using a panel of six isogenic *E. coli* clones expressing representative members of each of the major MCR groups (MCR-1, −2, −3, −4, −5, −8) and an *E. coli* clone carrying the expression vector (pDM1) alone (Table 1). For the *E. coli* clone carrying only the expression vector, the negative mass spectrum, scanned between *m/z* 1,600 and 2,200, is dominated by a set of peaks assigned to bis-phosphorylated hexa-acyl Lipid A. The major peak at *m/z* 1796.2 corresponds to hexa-acyl diphosphoryl Lipid A containing four C14:0 3-OH, one C14:0 and one C12:0, referred to as native Lipid A (Figure 1, top row). For *E. coli* clones expressing MCR enzymes, the addition of pETN to the 1-phosphate of native Lipid A leads to an additional peak (*m/z* 1919.2) shifted by +123 *m/z* compared to the mass of the major peak at *m/z* 1796.2 (Figure 1, bottom row). The sample optimization process aimed to achieve a higher than 10-fold signal to noise ratio for the peaks at *m/z* 1796.2 and *m/z* 1919.2. For this purpose, the sample preparation procedure was divided into three steps: *i*) acid hydrolysis, *ii*) sample washing and *iii*) sample resuspension prior to MALDI-TOF analysis. Parameters such as the acetic acid concentration, the time of hydrolysis, the sample washing procedure after acid hydrolysis and the sample density after resuspension were adjusted accordingly. The final optimized protocol is detailed in the Materials and Methods section.

**Figure 1:**
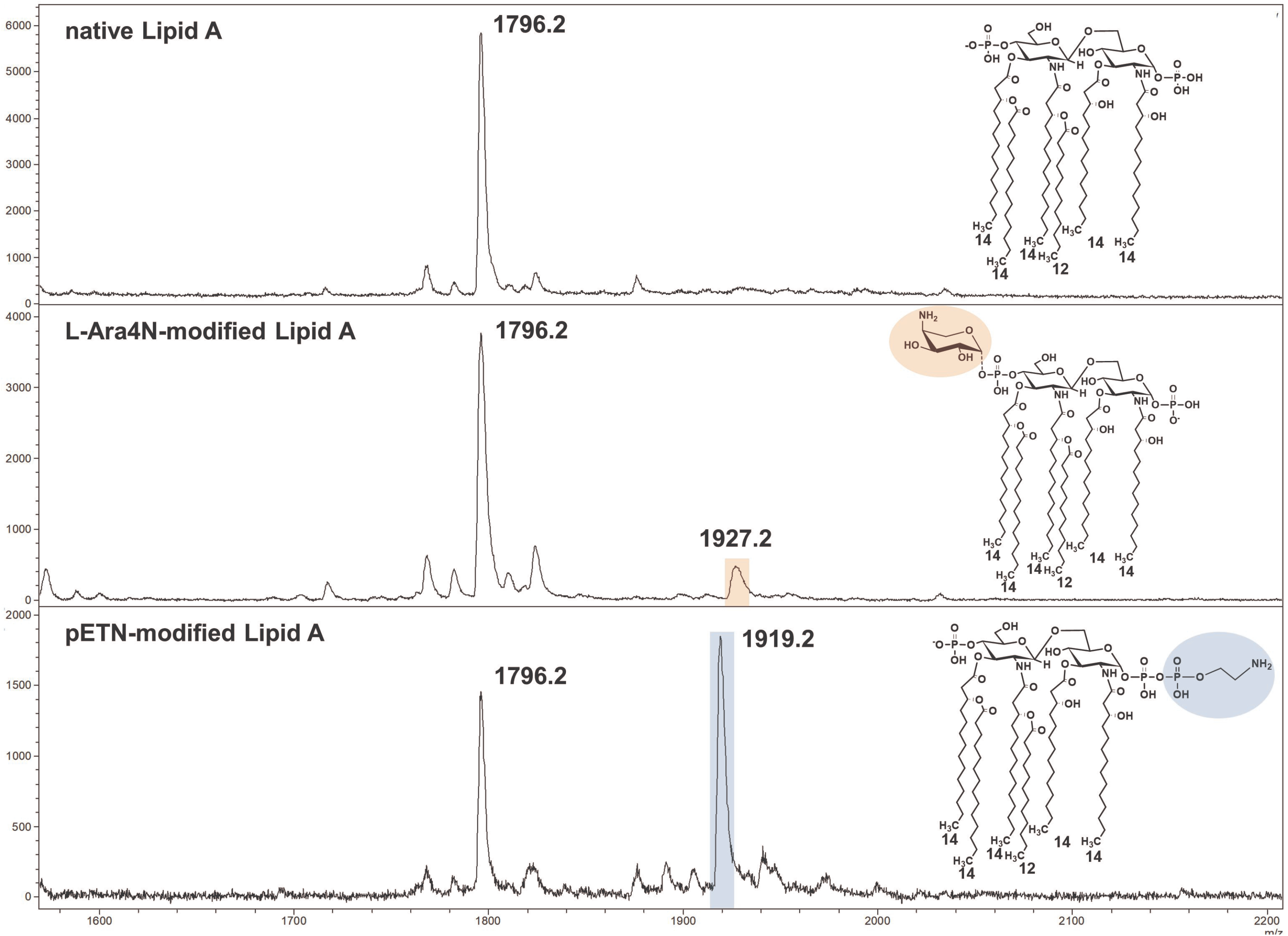
Representative mass spectra of native and modified *E. coli* Lipid A acquired using the linear negative-ion mode of a MALDI Biotyper Sirius system (Bruker Daltonics). Native *E. coli* Lipid A is detected as one major peak at *m/z* 1796.2 (top row). Lipid A from colistin-resistant *E. coli* isolates carrying chromosomal mutations is modified by L-Ara4N which is detected as an additional peak at *m/z* 1927.2 (highlighted in orange) (middle row) and/or pETN which is detected as an additional peak at *m/z* 1919.2 (highlighted in blue) (bottom row). Lipid A from strains exhibiting MCR-mediated resistance to colistin is only modified by pETN (bottom row); the spectrum shown is typical of an mcr-carrying isolate. Insets show the corresponding structures of native and modified Lipid A with the L-Ara4N and pETN modifications highlighted as appropriate.

The optimized version of the MALDIxin test was validated using a panel of 40 *E. coli* clinical isolates (Table 1), including 19 MCR producers, 12 chromosomally-resistant isolates and 9 colistin susceptible isolates. For all MCR producers in this panel both the native Lipid A peak and the additional pETN peak at *m/z* 1919.2 was observed, independent of the amino acid sequence of the MCR protein conferring colistin resistance. For colistin resistant isolates that were not found to carry an *mcr* gene by multiplex PCR (15), and are thus likely to harbor chromosomal mutations that lead to colistin resistance, we were able to detect a peak at *m/z* 1927.2, in addition to the native Lipid A peak. This signal corresponds to the addition of L-Ara4N to the 4’-phosphate of Lipid A, resulting in an increase of +131 *m/z* compared to the native Lipid A peak (Figure 1, middle row). In several of these isolates, peaks at both *m/z* 1919.2 and *m/z* 1927.2 were observed, suggesting that these organisms possess Lipid A species modified with both pETN and L-Ara4N (Table 1). Finally, for colistin susceptible *E. coli* clinical isolates, a single peak at *m/z* 1796.2 was detected, confirming that the Lipid A in these strains is unmodified. Using these spectra, PRR values for all strains were calculated. Susceptible *E. coli* strains have a PRR value of 0, whilst all colistin-resistant isolates have a positive PRR value (Table 1). Whilst this PRR value should be used to determine if an isolate is resistant or susceptible to colistin, the contribution of each Lipid A modification (Figure 1) to the overall PRR value, and thus colistin resistance, can be assessed by calculation of PRR^1919^ (pETN) and PRR^1927^ (L-Ara4N) values (Table 1).

## DISCUSSION

The work presented here broadens the applicability of our previously developed MALDIxin test (13) and represents an unbiased, fast, robust, cost-effective and high-throughput method to detect colistin resistance in *E. coli* by directly assessing the biochemical cause of resistance *i. e.* the modification of Lipid A. Therefore, unlike PCR-based testing, this method will reliably identify clinical isolates harboring chromosomal mutations, *mcr* genes and novel colistin resistance determinants, such as emerging MCR members, regardless of the genetic basis of resistance. Indeed, by determining the Lipid A modification(s) responsible for colistin resistance through the calculation of PRR^1919^ and PRR^1927^ values, potential MCR-producers (i.e. those organisms where the PRR value arises solely from the addition of pETN to Lipid A) can be identified for future in-depth characterization.

For this analysis we used the recently released MALDI Biotyper Sirius mass spectrometer. This system differs from previous Biotyper systems as it can operate in both positive and negative ion modes. Analytes that are acidic in nature, such as those containing phosphate or carboxylate groups, are more efficiently ionized by the generation of anions (16). As such, detection of Lipid A, which contains both long chain fatty acid and phosphate groups (at carbon 1 and 4’), is superior when anions are generated using the negative ion mode. Therefore, the newly introduced negative ion mode of the MALDI Biotyper Sirius allows efficient detection of both native Lipid A and its modified forms. Nonetheless, although the MALDI Biotyper Sirius is the optimal mass spectrometer for the assay as described here, Lipid A can also be detected using any MALDI-TOF mass spectrometer supporting negative-ion mode. In addition to the newly introduced negative ion mode, the MALDIxin test uses a super-DHB MALDI matrix, as opposed to the α-cyano-4-hydroxycinnamic acid (HCCA) matrix routinely used for bacterial identification by MALDI-TOF mass spectrometry. Whilst both super-DHB and HCCA are traditional organic matrices, super-DHB is a binary mixture of two benzoic acid derivatives. Mixed matrices such as super-DHB offer improved yields and signal-to-noise ratios for analyte ions by altering the co-crystallization of the analyte and matrix components (17). Together, these two advances allow the MALDI Biotyper Sirius to be used for both bacterial identification and robust colistin resistance determination, through detection of native or modified Lipid A from whole bacterial colonies.

The modification of Lipid A is a common mechanism of colistin resistance in organisms beyond *E. coli*. As the structure of Lipid A from a range of bacterial species (including *Klebsiella pneumonia, Shigella* spp. and *Pseudomonas aeruginosa*) can be determined by MALDI-TOF mass spectrometry (18), this technique provides a broadly applicable basis for the development of new diagnostics in many species of Gram-negative bacteria. Indeed, the Lipid A of *Salmonella* spp., which have been reported to carry MCR-enzymes (19) is similar to that of *E. coli* and can be detected using the negative-ion mode of the MALDI Biotyper Sirius as a peak at *m/z* 1796.2 (data not shown). Thus, it is likely that a similar +123 *m/z* addition to the native Lipid A peak will be observed in colistin resistant isolates of this organism. Similarly, Lipid A from *Acinetobacter baumannii* can be directly detected using MALDI-TOF mass spectrometry. Colistin resistance in this organism, primarily resulting from the overexpression of the chromosomally-encoded pETN transferase PmrC, can be detected as a +123 *m/z* addition to the peak corresponding to native bis-phosphorylated hepta-acyl Lipid A (13). These observations suggest that the optimized version of the MALDIxin test presented here will have broad utility in detecting colistin resistance in a range of Gram-negative bacteria.

The diagnostic assay described in this study will initially be made available to users of the MALDI Biotyper Sirius, along with full application support, for research use only (RUO) validation studies. This will be followed by the transformation of an already existing, RUO, web-based automated algorithm (Bruker Daltonics) into a new MALDI Biotyper software module. Dedicated MALDIXin consumables (e.g. pre-portioned purified matrix, calibration standards) will also be developed to enable simplified and standardized performance of the assay. The successful deployment of the new software module, in conjunction with MALDIxin specific laboratory consumables, will allow the subsequent introduction of *in vitro* diagnostic (IVD) consumables and software, following further clinical and analytical studies. These steps will ultimately bring the MALDIxin test into clinical laboratories in the near future.

Overall, this study represents a major step towards for the routine application of MALDI-TOF-based detection of colistin resistance and lays the foundations for a rapid diagnostic test for colistin resistance that will be readily accessible to most clinical microbiology laboratories. As such adoption of the MALDI Biotyper Sirius, and the subsequent introduction of the MALDIxin test, will facilitate improved management and treatment of patients with challenging MDR Gram-negative infections.

## ACKNOWLEDGMENTS

We acknowledge Prof. Youri GLUPCZYNSKI, Dr. Pierre BOGAERTS, Prof. Richard BONNET and IHMA Inc. Schaumburg for providing *E. coli* clinical isolates. This study was supported by the MRC Confidence in Concept Fund and the ISSF Wellcome Trust Grant 105603/Z/14/Z (to G.L-M), as well as the MRC Career Development Award MR/M009505/1 (to D.A.I.M.). L.D, A.F and G. L-M are co-inventors of the MALDIxin test for which a patent has been filed by Imperial Innovations.

